# Dietary macronutrients influence the gut microbiome and FA1PI, a non-invasive marker for GI health, in zoo-managed maned wolves (*Chrysocyon brachyurus*)

**DOI:** 10.64898/2026.08.10.743908

**Authors:** Morgan Bragg, Carly R. Muletz-Wolz, Elizabeth W. Freeman, Nucharin Songsasen

## Abstract

The maned wolf (*Chrysocyon brachyurus*) is a near-threatened canid species that suffers from gastrointestinal (GI) disease under human care. While the cause remains poorly understood, recent studies report that altered gut microbiota is linked to GI disease in the domestic dog. The goal of this study was to describe the relationship between gut bacteria and environmental factors, GI health, short chain fatty acid (SCFA) peaks, and genetic relatedness in maned wolves. Fresh fecal samples were collected twice from wolves (first collection n = 29; second collection n = 25) housed in six facilities in the United States. Fecal alpha-1 proteinase inhibitor (FA1PI), a biomarker for GI health in canids, and SCFA peaks were quantified. Information on diet, housing group and history of GI disease or inflammation was provided by the zoological facility for each wolf. We characterized gut microbiota using 16S rRNA gene amplicon sequencing with qPCR abundance corrections. We found that dietary fiber gross energy (GE), protein GE, and number of daily produce items influenced gut microbiota. Additionally, there was a negative relationship between dietary fiber GE and *Fusobacterium* sp. abundance, corroborating previous findings linking fiber to altered gut microbiome composition in canids. Lastly, fiber GE was negatively correlated with FA1PI, connecting fiber to GI health in the maned wolf. Findings from this study suggest that increasing dietary fiber GE changes the gut microbiome and may improve GI health in the maned wolf, highlighting important management decisions that can be considered to maintain a healthy zoo-managed maned wolf population.

**IMPORTANCE:** Findings from this study are important because gastrointestinal disease is a common health concern for maned wolves under human care, and the causes are not well understood. The current study found that diet, especially the amount of fiber gross energy, can affect gut bacteria and may support better gastrointestinal health. Diet alterations could be a practical tool for zoos to support management of gastrointestinal health issues in maned wolves. These findings also show the importance of carefully managing diets for a near-threatened species, since better gastrointestinal health may improve overall health and well-being and support reproduction. More research is needed to determine the best amount and type of fiber, but our results provide foundational information for improving care and supporting healthy zoo-managed maned wolf populations.

## 1. Introduction

The maned wolf (*Chrysocyon brachyurus*) is a near-threatened canid endemic to the grasslands of South America, with most individuals residing in the Brazilian Cerrado biome (Songsasen and Rodden 2010). Unfortunately, habitat loss and fragmentation associated with large scale agriculture production has reduced Cerrado vegetation by more than 50% with most of this area being unprotected (Pompeu et al. 2024), highlighting the importance of maintaining a healthy *ex-situ* population of maned wolves as a safeguard against extinction. While the zoo-managed population is integral for maned wolf conservation, it also is associated with health disorders not described in the wild. Specifically, gastrointestinal (GI) disease is a common cause of death in the US zoo population (Marcinczyk et al. 2024; Henson et al. 2017; Songsasen and Rodden 2010; Maia and Gouveia 2002). GI disease is defined as a group of diseases that are characterized by inflammation in the small or large intestinal mucosa and are frequently the cause of chronic vomiting and diarrhea in canids (Xu et al. 2016). While we do not know the underlying cause of GI disease in maned wolves, it is theorized that the cause of GI disease in domestic dogs is due to an immoderate immune system response triggered by factors like diet or commensal gut microbiota in individuals with a genetic predisposition (Dupouy-Manescau et al. 2024; Allenspach and Mochel 2022; Nishida et al. 2017; Marchesi et al. 2016; Suchodolski et al. 2012).

Gut microbiota provide their host with several benefits like supporting immune system development, synthesis of vitamins B and K, protection against pathogens, fermentation of non-digestible dietary carbohydrates and subsequent production of short-chain fatty acids (SCFAs; Jandhyala et al. 2015). Several factors, including diet and surrounding environment, can affect gut microbial community composition in wildlife species maintained in human care (Moustafa et al. 2021; Bragg et al. 2020; Cabana et al. 2019; Antwis et al. 2018; Grieneisen et al. 2017; McKenzie et al. 2017; Wasimuddin et al. 2017; Tung et al. 2015; Song et al. 2013). The maned wolf is a generalist omnivore and in the wild, their diet is composed of an equal ratio of small mammals and fruits/plant material (Paula and DeMatteo 2015). In the North American population, the maned wolf diet is made up of dry kibble, whole prey items and a variety of fruits and vegetables (Association of Zoos and Aquariums Canid Taxonomic Advisory Group 2012); however, the amount, brand and/or type of all food items can differ depending on zoological institution. Studies have shown that diet can impact gut microbiome composition in domestic dogs (Kim et al. 2017; Deng and Swanson 2015; Hang et al. 2012), red wolves (Bragg et al. 2020) and gray wolves (Chen et al. 2022); however, this relationship has not been explored in the maned wolf.

Similarly, modified social groups under human care could influence gut microbiome composition. In the wild, the maned wolf is generally monogamous with breeding pairs sharing resources and territory during breeding season and if juveniles are involved (Emmons 2012; Silveira et al. 2009). However, social behavior of the species can vary from solitary to more cohesive behavior between the pair depending on the individuals, environment and/or reproductive status (Emmons 2012; Dietz 1984). Previous research has provided evidence that co-habitation or group membership can influence gut microbiome composition in mammals (Antwis et al. 2018; Grieneisen et al. 2017; Tung et al. 2015; Song et al. 2013). In addition, Jones et al. (2018) found that female maned wolves housed with other females had significantly increased fecal glucocorticoid metabolite concentrations, a biomarker of stress, compared to female maned wolves house singly or with a male. It is possible that social stress in individuals could trigger a stress response, subsequently altering gut microbiome composition (Vlčková et al. 2018; Bailey et al. 2011; Galley et al. 2014) and providing opportunistic pathogens with favorable circumstances for colonization (Bailey et al. 2010), leading to poor GI health. Additionally, related individuals tend to have more similar microbiome composition (Tavalire et al. 2021; Wasimuddin et al. 2017) and this similarity, including both commensal and pathogenic taxa, could be vertically transmitted from mother to offspring (Schmidt et al. 2019; Ferretti et al. 2018; Moeller et al. 2018; Galley et al. 2014). It is beneficial to elucidate if and how co-habitation and genetic relatedness alter gut microbiome composition in zoo-managed maned wolves as it could predispose individuals to gut dysbiosis and poor GI health.

Dysbiosis in gut microbiota composition can lead to changes in SCFA production and concentration, leaving individuals at risk for poor GI health. The three main SCFAs produced in the gut are butyrate, acetate and propionate (den Besten et al. 2013; Ríos-Covián et al. 2016). These three SCFAs act as an energy source for epithelial cell growth and bacterial metabolism (Omori et al. 2017), moderate innate and adaptive immune cell production, movement and function (Gonçalves et al. 2018) and may relate to gut health and integrity. Butyrate, produced by bacteria in the phylum Bacteroidetes and Firmicutes, can suppress inflammation in immune and epithelial cells (Gonçalves et al. 2018). Acetate, produced by the phylum Bacteroidetes, is the main component that allows certain bacteria to kill pathogens within the GI tract (Ríos-Covián et al. 2016). Lastly, propionate is also produced by the phylum Bacteroidetes and has anti-inflammatory effects (Ríos-Covián et al. 2016). As a result of their influential mechanisms, any alteration in presence or concentration of these SCFAs can have negative impacts on GI health.

The microbiome is a complex environment, and external stressors can influence gut microbiome composition in either a stochastic or deterministic manner. The response to these stressors can vary by individual or species (Zaneveld et al. 2017), making it challenging to identify general patterns associated with GI health. Assessment of how external factors influence gut microbiome composition over time can provide information about the stability of the gut microbiome, which could impact GI health. The objective of this study was to examine the relationship between the gut microbiome, environmental factors, SCFA composition, genetic relatedness and GI health in zoo-managed maned wolves and assess if patterns identified remain consistent over time. Findings from this study will allow us to better understand the influence of management choices on the gut microbiome and GI health in zoo-managed maned wolves.

## 2. Methods

### 2.1 Animals and sample collection

We collected fecal samples from a total of 30 wolves across six different zoological facilities. There were 14 males and 16 females that were ages four to 12 (Table 1). Fresh fecal samples obtained within one hour of defecation were collected in two rounds for this study. The first collection took place between August and November 2019 and included 29 maned wolves. The second fecal collection was completed six months later, between January and May of 2020, and included 25 of the 29 wolves from the first collection with the addition of one new wolf. The four wolves from the first collection that were not included in the second collection was because of institution transfer or euthanasia for reasons that were unrelated to this study. Fecal samples were collected opportunistically during routine husbandry for three consecutive days by staff at each zoological facility and kept at −20°C until shipment to Smithsonian National Zoo & Conservation Biology Institute.

**Table 1.**
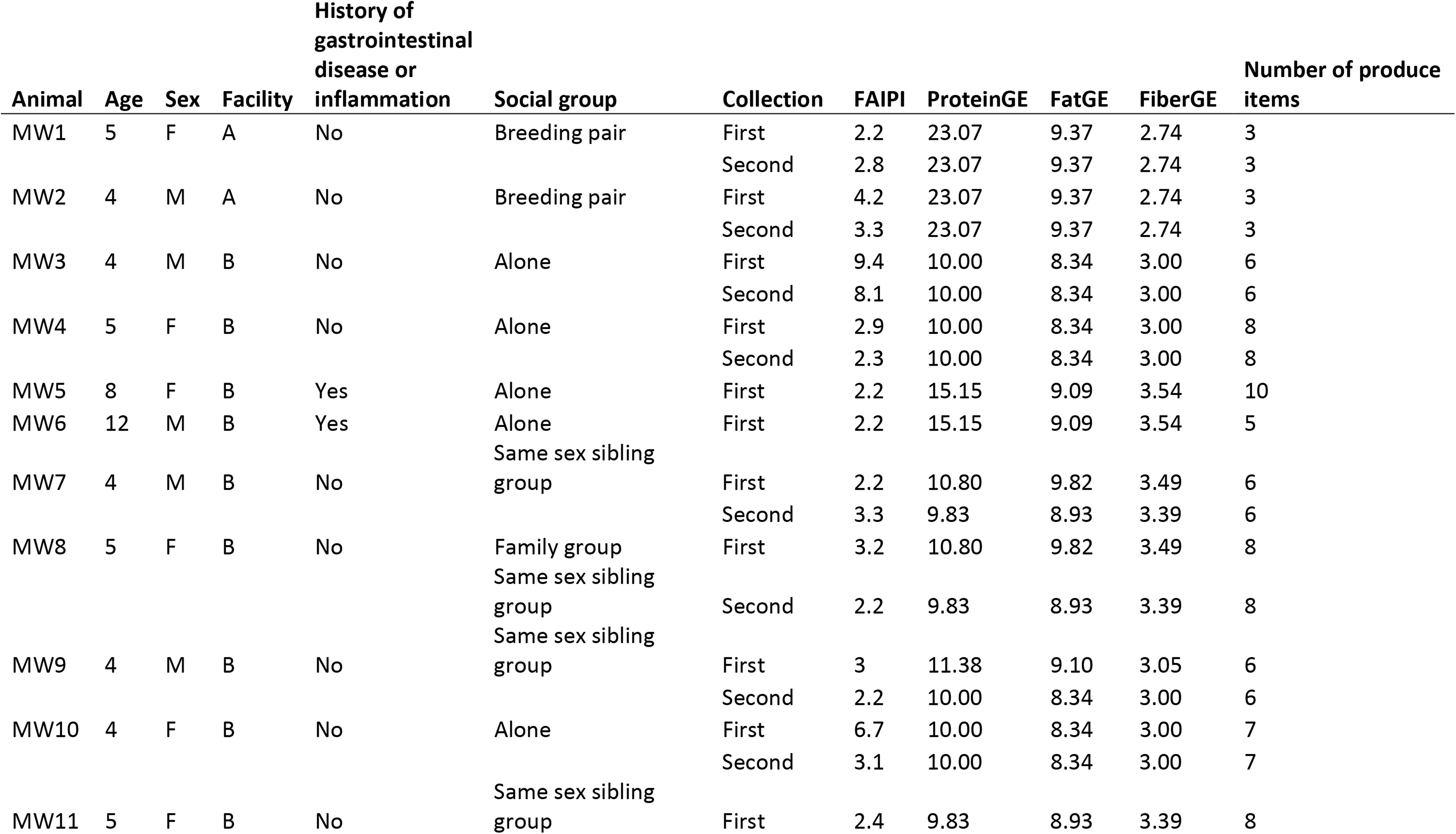
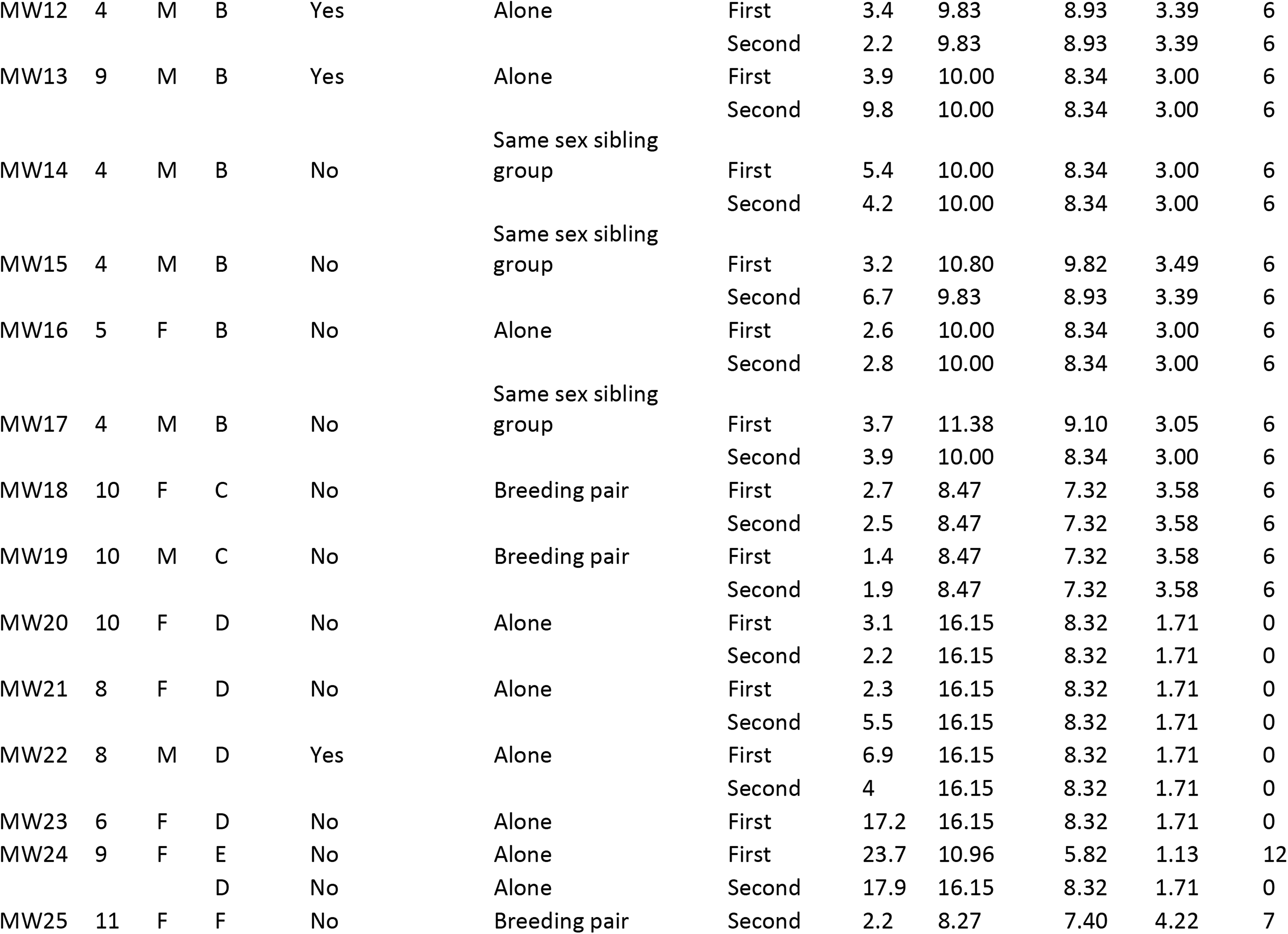
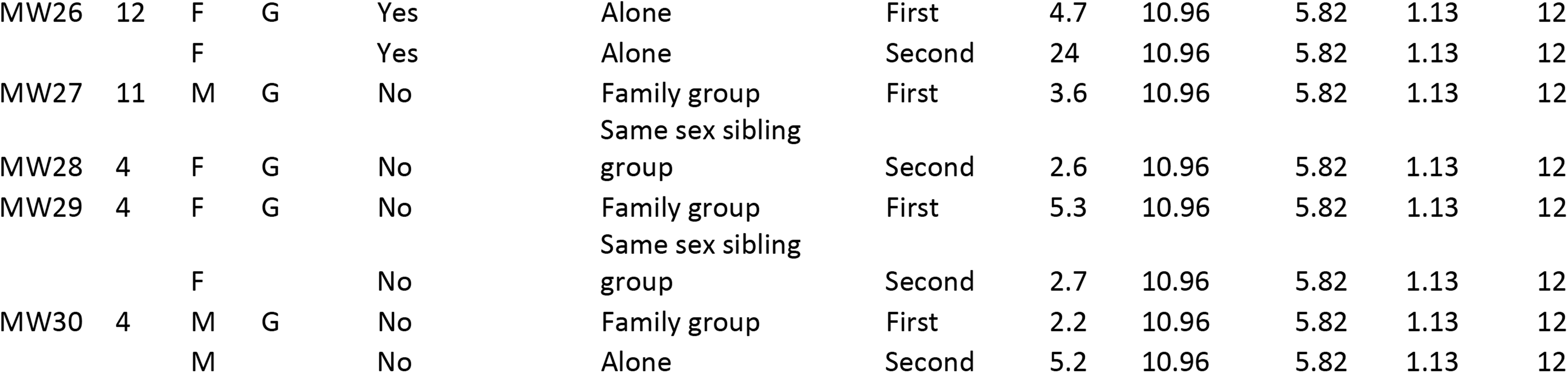
List of 30 participating maned wolves and their age, sex, facility, history of gastrointestinal (GI) disease or inflammation, number of produce items in their daily diet, composition of social group, collection participation, fecal alpha-1 proteinase inhibitor (FA1PI) concentrations and daily diet protein, fat and fiber gross energy (GE) amount for each wolf.

### 2.2 Animal management

Maned wolves were maintained alone, as a breeding pair, in a same sex sibling group or in a family group.

Wolves were fed their standard daily diet that includes dry kibble (meat-based dry kibble for domestic dogs [Natural Balance or Hill’s Science Diet] or dry kibble created specifically for maned wolves [Mazuri]), a variety of protein items (i.e., jumbo rats, Milliken brand small carnivore diet, mice, chicks, herring, hard boiled eggs, etc.) and produce (i.e., banana, papaya, sweet potato, apple, tomato, etc.). The amount, type and/or brand of dietary components varied across zoological institutions.

### 2.3 Diet analysis

Animal keepers at each institute provided daily diet information for participating wolves, which included the amount, brand (if applicable), variety and frequency of kibble, produce and protein/whole prey items offered. Due to the high variability in the diet offered to wolves among institutions, each daily diet was broken down into total energy (total kcals), protein (%DM), fiber (%DM), fat (%DM) and moisture (%) using Zootrition® Dietary management database (© E. Dierenfeld, St. Louis, MO, 2011). Then, protein (%DM), fiber (%DM), fat (%DM) were converted to gross energy (GE) to avoid performing statistics on percentages. Gross energy (mg/kcal) for each nutrient category was calculated by dividing the percent of each nutrient category by the total energy amount then multiplying by 1,000 (Keady et al. 2023). From here forward, we refer to the nutrient mg/kcal GE as protein GE, fiber GE and fat GE.

### 2.4 Assessment of gastrointestinal health

The gold standard for diagnosing GI disease includes performing an endoscopy to assess mucosal surface of the duodenum and taking full-thickness biopsies to analyze histology (Wolf 2019; Seeley et al. 2016). Specialized equipment and expertise are required for this, which are not easily accessible for most participating facilities. Therefore, we used fecal alpha 1-proteinase inhibitor (FA1PI), a non-invasive marker for protein losing enteropathy in human (Thomas et al. 1981), domestic dog (Murphy et al. 2003), and domestic cats (Burke et al. 2013) as a proxy for GI health. A subsample was taken from each fecal sample and sent to Texas A&M Gastrointestinal Laboratory for FA1PI quantification. In adult domestic dogs, the standard to assess abnormal GI health status is a three-day FA1PI mean concentration ≥ 13.9 μg/g or one individual sample ≥ 21μg/g (Texas A&M Gastrointestinal Laboratory 2021). However, we do not know what FA1PI values in maned wolves indicate abnormality. Generally in domestic cats and dogs, individuals with normal GI health status have low concentrations of FA1PI compared to individuals with GI disease (Hibbetts et al. 1999; Murphy et al. 2003; Burke et al. 2013). Therefore, we chose to examine the relationship between environmental factors and FA1PI, and if we observed a linear relationship, we suggested that specific environmental factor is linked to GI health. In addition to FA1PI analysis, previous history of GI disease or inflammation (yes/no) for each wolf were obtained from animal keepers at each institution. For this study, history of GI disease or inflammation was defined as being clinically treated for GI disease within three years prior to fecal sample collection or GI tract inflammation present during post-mortem necropsy.

### 2.5 Short-chain fatty acid analysis

An additional subsample was taken from one fecal sample (Day 1, first collection) from 29 out of the 30 wolves and sent to George Mason University Metabolomics Core (Manassas, VA) for SCFA analysis via solid phase microextraction in conjunction with gas chromatography – mass spectrometry (GC-MS) and gas chromatography – flame ionized detection (GC-FID; Dixon et al. 2011). Peak heights were obtained for acetic acid, propanoic acid, butanoic acid, pentanoic acid, hexanoic acid and octanoic acid as these are common metabolites associated with GI health in canids (Higueras et al. 2024; Sun et al. 2023; Jewell et al. 2022).

### 2.6 Microbiome analysis

Protocol for microbiome extraction and library preparation followed methods published in Bragg et al. (2020). Briefly, DNA was extracted using 0.25g of thawed, wet fecal sample and the QIAamp PowerFecal Pro kit (Cat. 51804, Qiagen). Negative and positive controls (Zymobiomics microbial community standard; Cat. D6300, Zymo) were included for each round of extraction. Library preparation entailed a two-step polymerase chain reaction (PCR) protocol combined with a dual-index paired-end Illumina sequencing on a MiSeq (v3 chemistry: 2 × 300 bp kit). We used universal primers 515F-Y (GTGYCAGCMGCCGCGGTAA) and 939R (CTTGTGCGGGCCCCCGTCAATTC) to target the V4-V5 region of the 16S rRNA gene. Negative and positive DNA controls (Zymobiomics microbial community DNA standard; Cat. D6305, Zymo) were included for each round of PCR.

Filtering low quality sequences, checking for chimeras and trimming of data was done using the R package “dada2” (Callahan et al. 2016). Amplicon sequence variants (ASVs) were generated, and taxonomy assigned by aligning the sequences against the Ribosomal Database Project (RDP) 16S training set 16/release11.5 (Wang et al 2007). The R package “decontam” (Davis et al. 2017) was used to identify contaminants using the Fisher method with a threshold of 0.1, which removed four ASVs. We filtered out singletons ASVs (ASV only occurs one sequence in one sample), negative controls, Cyanobacteria and mitochondria. The bacterial taxa we expected to be in the positive controls were found and removed prior to analyses. The variation in sequencing depth was approximately 7x (min = 8,010, max = 54,706) across individuals; therefore, we did not implore rarefying techniques following recommendations by Weiss et al. (2017).

### 2.7 Quantitative PCR

We normalized sample-wise differences in total bacterial abundance and estimated the absolute abundance of each ASV by conducting quantitative PCR (qPCR) using SsoAdvanced Universal SYBR Green Supermix (Cat. 1725270; Bio-Rad). We input 30ng (10ng/ul) of DNA into each reaction and samples were run in duplicates. The same primers used for sequencing, 515F-Y and 939R, were utilized. Amplification was carried out with a 3-minute denaturing step at 95°C followed by 40 cycles of 15 second denaturation at 98°C and 45 second annealing at 60°C. Gblock standards were created following Zemb et al. (2020) and ran in duplication at concentrations of 10^9^, 10^8^, 10^7^, 10^6^, 10^5^, 10^4^, 10^3^ along with each set of reactions as well as duplicate PCR negative controls. The Cq values from each duplicate were averaged and divided by 10 to quantify the number of 16S rRNA copies per ng of DNA. Absolute abundance was estimated by multiplying the relative abundance of each ASV by the number of 16S rRNA copies per ng of DNA for each sample (Epp Schmidt et al. 2022; Jian et al. 2018).

### 2.8 Statistical analysis

We conducted all statistical analyses in R (version 4.1.2; R Core Team 2021) and significance was determined as p value ≤ 0.05. Prior to analyses, we subset the dataset to only include the sample collected on Day 1 from both collections.

Pearson correlation coefficient (function *cor*) was calculated among dietary variables (protein GE, fiber GE, fat GE, number of produce items) and variables with coefficients above 0.5 were considered correlated. High correlation prevented dietary explanatory variables (protein GE x number of produce items r = −0.55, fat GE x number of produce items r = −0.61, fat GE x fiber GE r = 0.67) from being included in the same model. To overcome this, we subset the dietary variables and conducted a principal component analysis (PCA; function *princomp*; Supplementary Table 1). Principal component (PC) 1 scores were extracted and added back to the full dataset prior to analyses. We refer to this variable as diet PC1 moving forward. Additionally, we performed a log10 transformation on FA1PI concentrations to improve normality of the variable and the transformed variable was used in all models hereafter.

Prior to analyses, we determined the appropriate random effect of facility, animal identity or animal identity nested within facility with restricted maximum likelihood (REML) on the full model and compared the models with Akaike’s information criterion (AIC) using the *step* function in the “lmerTest” package (Kuznetsova et al. 2017). Those results indicated that animal identity nested inside of facility was the appropriate random effect to account for repeated sampling from the same individuals at different zoological facilities. Therefore, animal identity nested inside of facility was used as a random effect for all alpha and beta diversity models.

#### 2.8.1 Comparison between first and second collection

To compare alpha diversity between the first date range and second date range collection, we ran linear mixed effect models on a merged data set containing the first and second collection with species richness or Faith’s phylogenetic diversity as the response variable and collection as the explanatory variables with animal identity nested inside of facility as the random effect.

To compare beta diversity between first date range and second date range collection, a permutational multivariate analysis of variance (PERMANOVA; function *adonis2*, “vegan” package; Oksanen et al. 2022) was used with Bray-Curtis, Jaccard’s or unweighted UniFrac distances as the response variable and collection and facility were the explanatory variables with animal identity as the random effect.

#### 2.8.2 Alpha diversity

Alpha diversity was measured using species richness and Faith’s phylogenetic diversity. Correlations among explanatory variables were tested using generalized variance inflation factors (VIF; function *vif*, “car” package; Fox and Weisburg 2019). Variables with a VIF value greater than five were considered correlated and excluded from being in the same model. To test these relationships, we used linear mixed effect models (LMMs, function *lmer*, “lme4” package; Bates et al. 2015) with species richness or Faith’s phylogenetic diversity as the response variable and age, sex, diet PC1, FA1PI concentration, social group and history of gastrointestinal disease or inflammation as explanatory variables with animal identity nested inside of facility as the random effect. We ran a corresponding null model for each linear mixed effect model. We used AIC to identify the best model.

#### 2.8.3 Beta diversity

To quantify the influence of environmental variables and GI health on microbiome composition, we normalized counts based on cumulative sum scaling prior to analysis (function cumNorm, “metagenomeSeq” package; Paulson et al. 2013) and PERMANOVAs (function *adonis2*, “vegan” package) were used with Bray-Curtis, Jaccard’s or unweighted UniFrac distances as the response variable and age, sex, social group, FA1PI, history of GI disease or inflammation, diet PC1 and facility as the explanatory variables with animal identity nested in facility as the random effect.

#### 2.8.4 Distance-based redundancy analysis

We used distance-based linear modeling (function *capscale*, “vegan” package) to assess the relationship between individual dietary variables (protein GE, fiber GE, fat GE, number of produce items) and gut microbiome composition. Models were run with Bray-Curtis, Jaccard and unweighted UniFrac distances. Final model selection was done using forward and backward model selection (function *ordiR2step*, package “vegan”).

We used additional distance-based linear models to assess the relationship between individual SCFA peak heights (acetic acid, propanoic acid, butanoic acid, pentanoic acid, hexanoic acid and octanoic acid) and gut microbiome composition. The dataset was subset to only include samples from wolves that had corresponding SCFA concentrations (n = 29; one wolf was accidentally not included in SCFA analysis). Models were run with Bray-Curtis, Jaccard and unweighted UniFrac distances. Final model selection was done using forward and backward model selection.

#### 2.8.5 Kinship values and gut microbiome composition

Pedigree information was available for 22 wolves in this study and kinship values were determined from pedigrees where 0 represented unrelated wolves, 0.125 for great-grandparents/grandchildren, 0.25 for grandparents/grandchildren, 0.5 for parent/offspring and for full siblings. We conducted a mantel test (function *mantel.test,* “ape” package; Paradis and Schliep 2019) to test the relationship between kinship values and microbiome composition.

#### 2.8.6 ASVs linked to fiber GE & SCFAs

Given the known relationship between fiber and SCFAs to GI health, we assessed relationships between individual bacterial strain abundance and fiber GE using linear mixed effects models. Taxa were filtered to only include ASVs with at least 1% abundance and that were present in a minimum of three samples (n = 128 ASVs). For fiber GE analysis, the response variable was log10 transformed bacterial abundances and fiber GE was the explanatory variables with animal nested inside of facility as the random effect. For SCFA analysis, propanoic, butanoic and pentanoic acid were the most abundant SCFAs quantified in our study and were the focus of our analysis with the response variable being log10 transformed bacterial abundances and propanoic, butanoic, or pentanoic acid as the explanatory variables with facility as the random effect. Taxa were filtered to only include ASVs with at least 0.1% abundance and that were present in a minimum of 16 samples (n = 20 ASVs). High spearman correlation values (r < 0.5) between propanoic acid and butanoic and pentatoic acid prevented all three SCFAs from being included in the same model. P values were adjusted for multiple comparisons with Benjamini and Hochberg correction.

#### 2.8.7 Structural equation model

In our analysis, we observed fiber GE showing correlations with both *Fusobacterium* sp. and FA1PI concentrations and wished to determine if these effects were direct or indirect. Therefore, we tested direct and indirect effects of fiber GE and *Fusobacterium* sp. abundance on log10 FA1PI concentrations using structural equation modeling (n = 53; function *psem*, package “piecewiseSEM”; Lefcheck 2016). We assessed two *a priori* models; one, a full model representing all possible relationships between variables and two, a model assuming no direct relationship between *Fusobacterium* sp. abundance and log10 FA1PI concentrations. Models were compared using ANOVA p-values and AIC. If no significant differences were detected or if AIC values fell within 2 points of each other, the most parsimonious model was interpreted as the final model.

## 3. Results

We obtained a total of 2,990,662 high quality bacterial sequences from 53 maned wolf samples (mean = 20,022 sequences, min = 6,756 sequences, max = 48,089 sequences). A total of 517 bacterial ASVs were identified from the phyla Firmicutes, Actinobacteria, Bacteroidetes, Fusobacteria, Proteobacteria, Chloroflexi and Campilobacterota, respectively. Across both the first and second collections, FA1PI concentrations ranged from 1.4 – 24 µg/g (median = 3.2 µg/g). Propanoic (median peak height = 7,106,149) and butanoic acid (median peak height = 6,907,639) were the most represented SCFAs (Supplementary Table 2).

No differences in species richness, Faith’s phylogenetic diversity or bacterial composition (p > 0.05) were seen between the first date range and second date range collection periods. Therefore, analyses were conducted using a dataset with the first date range and second date range collection periods compiled together with animal identity as a random effect to account for repeated sampling of the same individuals.

### 3.1 Microbiome structure: alpha diversity

At both collection periods, there was no relationship between alpha diversity measures of the gut microbiome (species richness and Faith’s phylogenetic diversity) and age, sex, diet PC1, social group, FA1PI or history of GI disease or inflammation (LMMs: p > 0.05).

### 3.2 Microbiome structure: beta diversity

At both collection periods, there was no relationship between beta diversity measures of the gut microbiome (Bray-Curtis, Jaccard, and unweighted UniFrac composition) and age, sex, diet PC1, social group, FA1PI or history of GI disease or inflammation, (PERMANOVA: p > 0.05). At both collection periods, no relationship was found between kinship values and microbiome composition (PERMANOVA: p > 0.05). At the first collection period, there was no relationship between SCFA profile and microbiome composition (PERMANOVA: p > 0.05).

### 3.3 Individual diet components, microbiome composition and GI health

We found that individual diet components fiber GE, fat GE, protein and number of produce items were correlated to linear changes in microbiome composition (Protein GE dbRDA: Jaccard p = 0.02, Unweighted UniFrac p = 0.04; Fiber GE dbRDA: Bray Curtis p = 0.002, Jaccard p = 0.001, unweighted UniFrac p = 0.003; Produce dbRDA: Bray Curtis p = 0.01, Jaccard p = 0.002, unweighted UniFrac p = 0.002; Figure 1). A relationship was documented between fat GE and microbiome composition only when using Jaccard distances (p = 0.02). When examining the relationship between individual diet components and GI health, we noted a negative relationship between fiber GE and FA1PI concentrations (LMM: p = 0.003, t_28.6_ = −3.29; Figure 2) but no influence of fat GE, protein GE or number of produce items on GI health (p > 0.05). Specifically, when fiber GE increased by one unit, the FA1PI concentration decreased by approximately 0.14 ng/µl.

**Figure 1.**
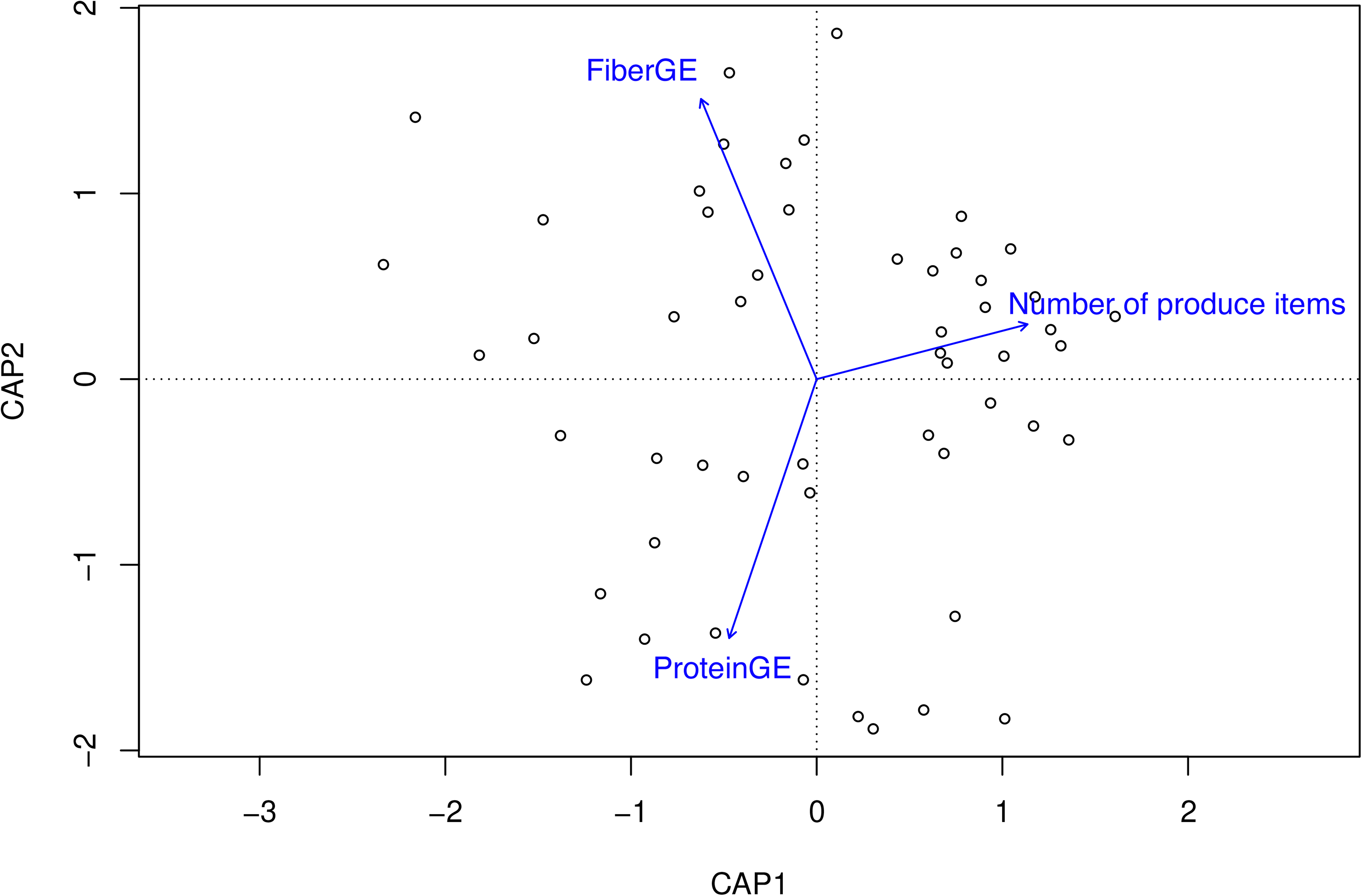
Gut bacteria composition changed in a linear manner with dietary fiber gross energy (GE), number of daily produce items and protein GE. Based on Unweighted UniFrac composition measure and distance-based linear models. Proportion of the total variance explained by constrained axes is 13.7%.

**Figure 2.**
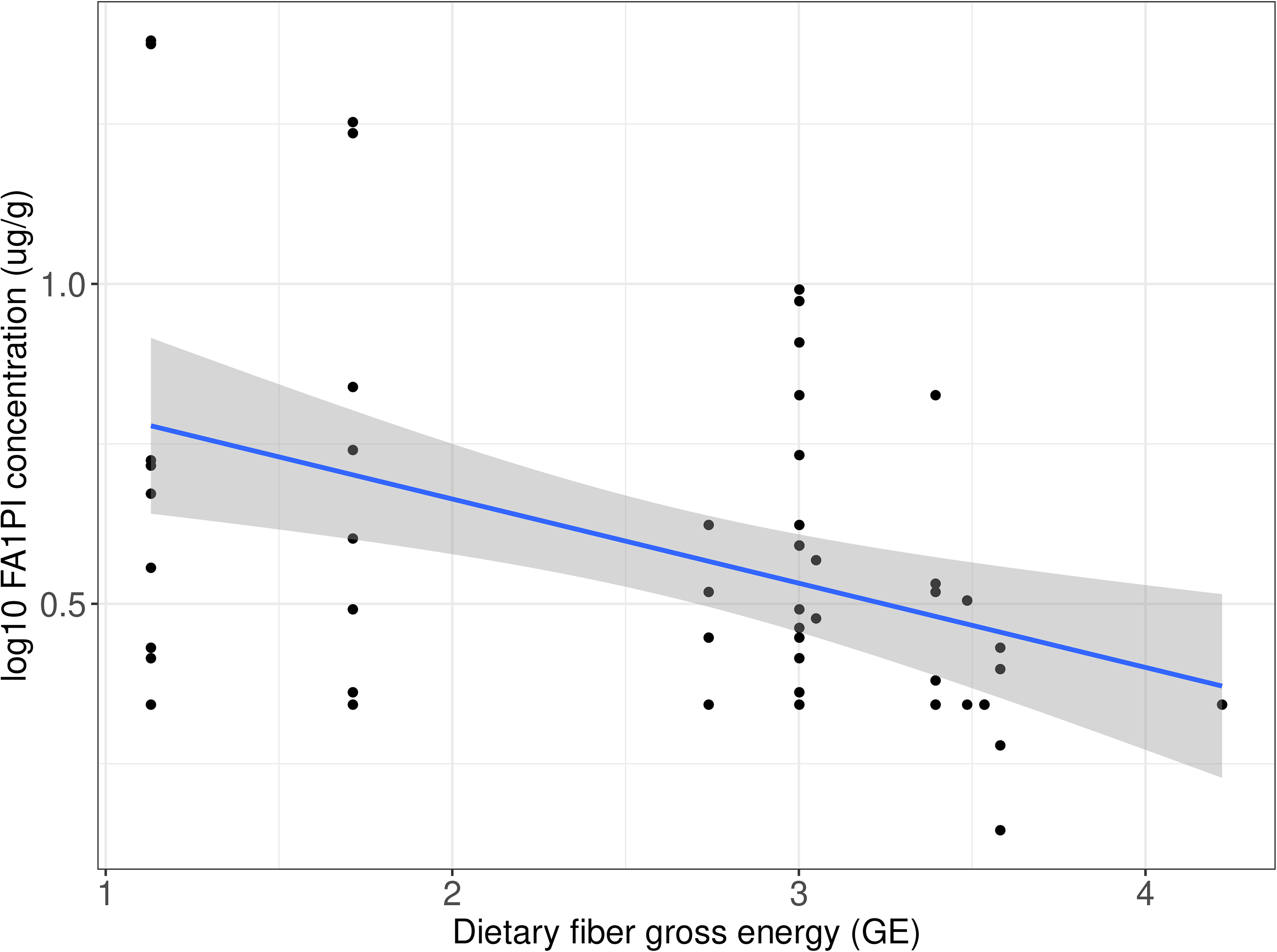
Negative relationship between dietary fiber gross energy and fecal alpha1-proteinase inhibitory (FA1PI) concentrations, a bio-marker for GI health (higher values indicating abnormal GI conditions), in zoo-managed maned wolves (p = 0.003).

### 3.4 ASVs linked to GI health and fiber GE

We discerned if the abundance of any bacterial ASVs (n = 128) were related to fiber GE. We detected one ASV (*Fusobacterium* sp. ASV25 p = 0.03) that had a negative relationship with fiber GE after adjusting for multiple comparisons (Figure 3).

**Figure 3.**
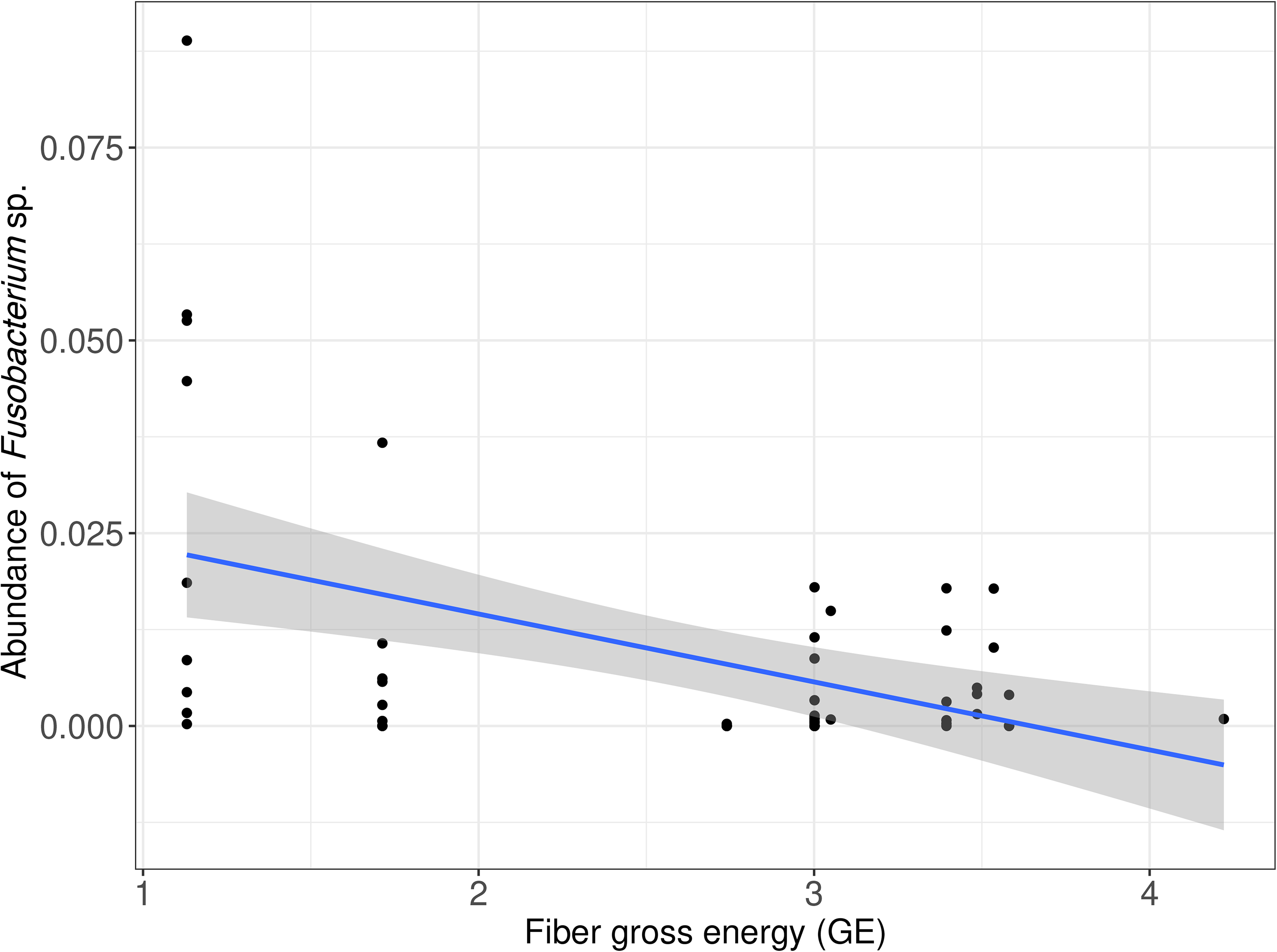
Negative relationship between abundance of *Fusobacterium* sp. (ASV25 p = 0.03) and fiber gross energy (GE).

### 3.5 ASVs linked to propionate, butanoic, and pentanoic acid

We discerned if any ASVs (n = 30) were related to propionate, butanoic, and pentanoic acid. One ASV, *Clostridium sensu stricto* sp. (ASV11 p > 0.001; Figure 4), had a negative relationship with propanoic acid. We identified 14 ASVs that were significantly associated with butanoic acid (*Blautia* sp, ASV22 featured in Figure 4; full list available in Supplementary Figure 1; p < 0.02). Further, we highlighted 28 ASVs that were significantly linked to pentanoic acid (*Megamonas* sp. ASV10 and *Blautia* sp. ASV15 featured in Figure 4; full list available in Supplementary Figure 2; p < 0.003).

**Figure 4.**
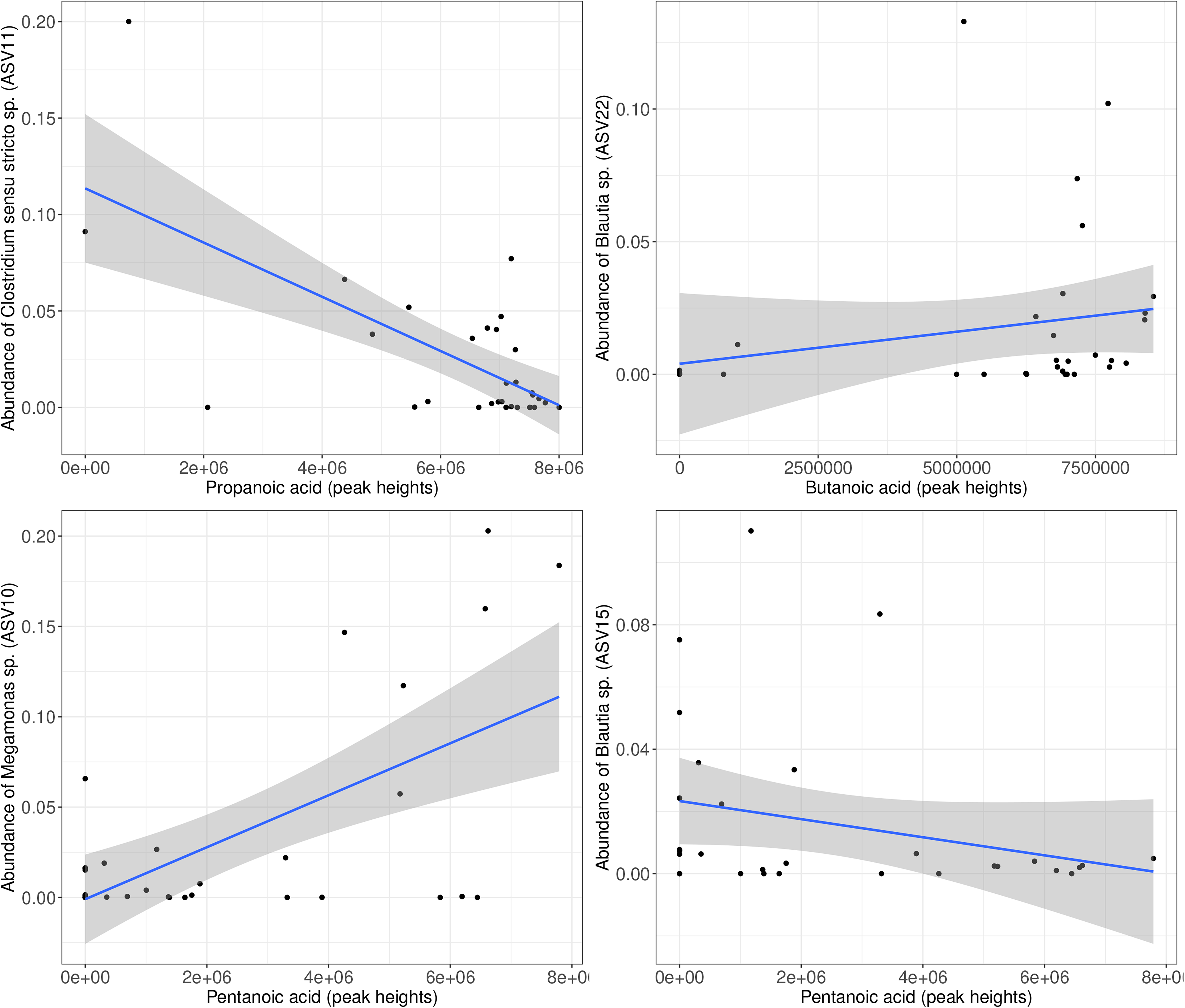
(A) Negative relationship between abundance of *Clostridium sensu stricto* sp. (ASV11; p < 0.001) and propanoic acid, (B) Positive relationship between abundance of *Blautia* sp. (ASV22; p < 0.001) and butanoic acid, (C) Positive relationship between *Megamonas* sp. (ASV10; p < 0.001) and pentanoic acid, and (D) Negative relationship between abundance of *Blautia* sp. (ASV15; p < 0.001) and pentanoic acid.

**Figure 5.**
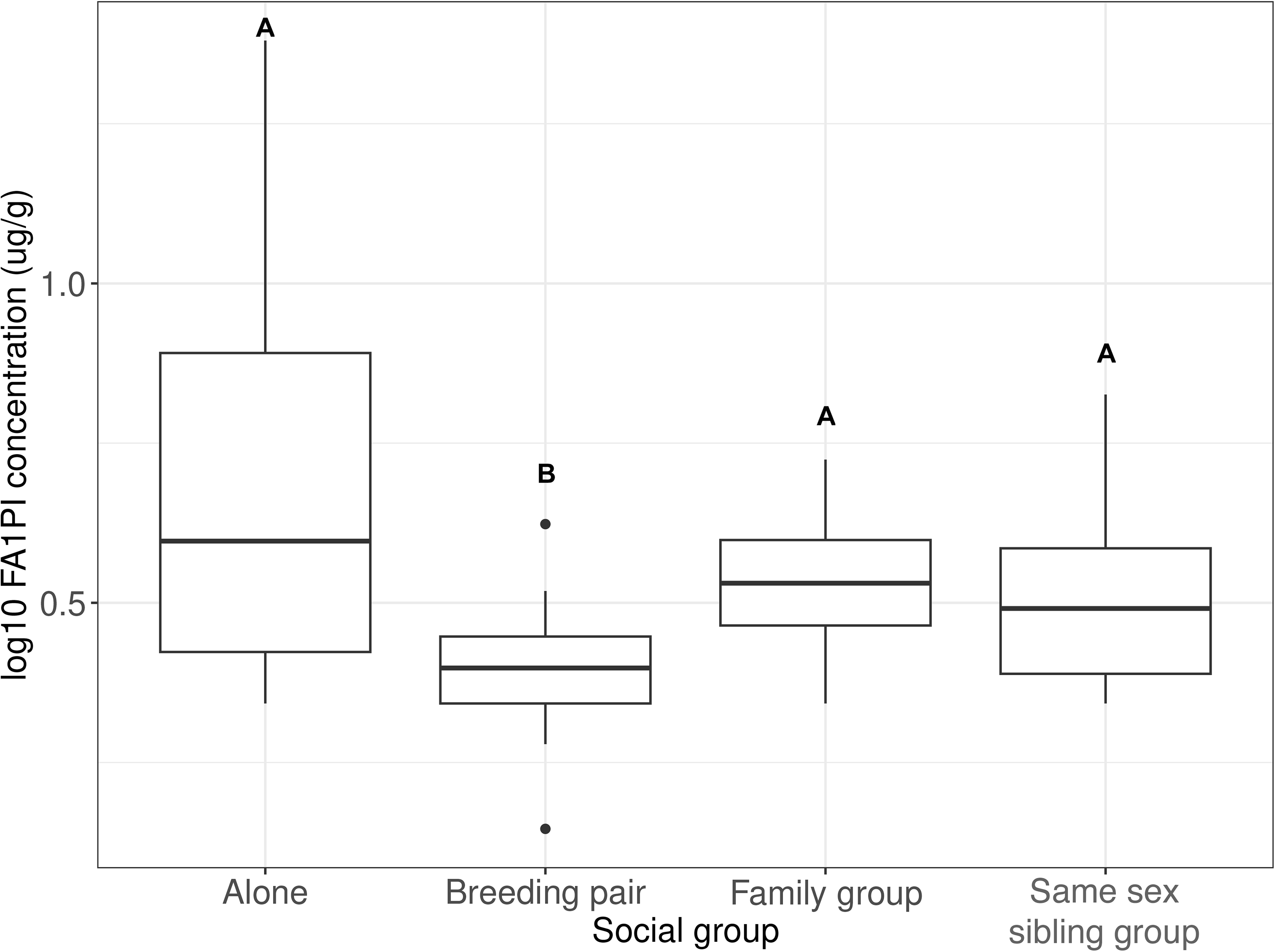
Fecal alpha1-proteinase inhibitory (FA1PI) concentrations among the four social groups zoo-managed maned wolves could live in; Wolves housed alone had higher concentrations of FA1PI compared to wolves housed as breeding pairs (p = 0.05)

### 3.6 Environmental variables and GI health

Social group influenced FA1PI concentrations, particularly maned wolves housed alone had higher FA1PI concentrations compared to wolves living in a breeding pair (LMM: p = 0.005, t_27.3_ = −3.06) and same sex sibling group (LMM: p = 0.02, t_36.7_ = −2.35). However, pairwise comparisons revealed only a significant difference between wolves housed alone compared to wolves housed as a breeding pair (estimate = 0.41, SE = 0.15, p = 0.05). No relationship between FA1PI, age, sex, diet PC1 or history of gastrointestinal disease or inflammation was observed (LMMs: p > 0.05).

### 3.7 Structural equation model

We found that fiber GE significantly influenced *Fusobacterium* sp. abundance (standardized coefficient = −0.46, p = 0.0005) and FA1PI concentrations (standardized coefficient = −0.43, p = 0.002), but there was no indirect influence of *Fusobacterium* sp. on FA1PI concentrations (Figure 6). This reflects the direct but separate influence of fiber GE on *Fusobacterium* sp. and FA1PI concentrations.

**Figure 6.**
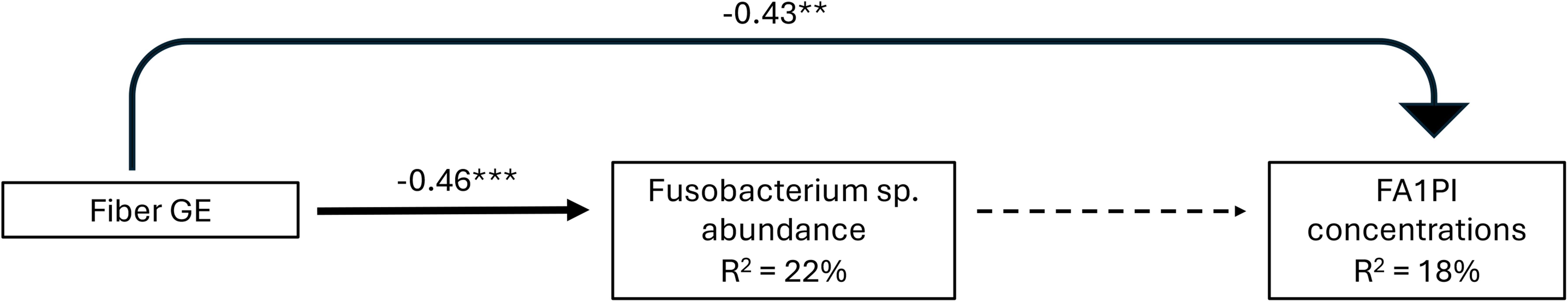
Direct and indirect effects of fiber gross energy (GE) and *Fusobacterium* sp. abundance on fecal alpha1-proteinase inhibitor (FA1PI) concentrations in zoo-managed maned wolves. Line thickness is scaled in proportion to the associated effect size, with dashed lines representing non-significant paths. *** represents p < 0.0001 and ** represents p < 0.01.

### 3.8 Kinship values and microbiome composition

There was no relationship (p > 0.05) between kinship values and microbiome composition found in this study.

## 4. Discussion

It is recognized that the gut microbiome plays an influential role in maintaining GI health in mammals. However, little information is known about gut microbiomes of maned wolves and how environmental factors impact the gut microbial community and GI health. We provided evidence that gut microbiome composition was stable over two time points six months apart, and dietary components like fat GE, fiber GE and the number of produce items influenced variation in gut microbiome composition. Specifically, fiber GE had separate and negative direct effects on FA1PI concentrations and *Fusobacterium* sp. abundance. Social group was also linked to FA1PI concentrations with wolves living alone having higher concentrations compared to wolves housed as a breeding pair or same sex sibling group. While we did not detect any environmental effects on overall SCFA composition, we did identify several ASVs linked to butanoic, pentanoic, and propanoic acid. Specifically, we highlighted (i) positive relationships between butanoic acid and abundance of *Blautia* sp. and pentanoic acid and abundance of *Megamonas* sp. and (ii) negative relationships between propanoic acid and abundance of *Clostridium sensu stricto* sp. and *Blautia* sp. and pentanoic acid. Findings from this study suggested that quantity of dietary macronutrients can influence gut microbiome composition and GI health in maned wolves. In addition, this study provides evidence linking specific bacterial taxa to individual SCFAs, suggesting the potential ability of these taxa to modulate GI health in the maned wolf.

It is well understood that diet can alter gut microbiomes in canids (Li et al. 2024; Martínez-López et al. 2021; Bragg et al. 2020; Kilburn et al. 2020; Schauf et al. 2018; Panasevich et al. 2013). Changes in diet can influence gut bacterial abundance and/or structure, which subsequently can alter metabolite production, immune system function and nutrient uptake (Moreno et al. 2022; Williams et al. 2019). In the present study, overall diet composition did not influence gut microbiome composition, but individual diet components like fat GE, fiber GE, and the number of produce items altered gut microbiome composition in a linear manner.

We found that fiber GE had a negative relationship with FA1PI concentrations, suggesting a potential mechanism to improve GI health by increasing fiber in maned wolf diet regimes. Dietary fiber is a critical nutrient required to support GI health via SCFA production, intestinal mucosal immunity, reduction of mucosa adherence, and reducing the number of pathogenic species (Cave et al. 2023). In this context, we hypothesized that there would be a negative relationship between fiber and FA1PI concentrations. In support of our hypothesis, we reported lower FA1PI concentrations, implying no or minimal GI mucosal inflammation, associated with higher fiber GE values in the current study. These results suggest there is an association between dietary fiber and GI health in the maned wolf. Although, it is important to note that we cannot infer nutrient digestibility and metabolized energy from GE values alone (Jha and Woyengo 2022; Cervantes-Pahm et al. 2013). Further research on dietary fiber digestibility and metabolized energy in relation to FA1PI concentrations in maned wolves is required to fully elucidate the link between fiber and GI health.

In the present study, we detected fiber GE having a negative relationship with abundance of *Fusobacterium* sp. in zoo-managed maned wolves. The genus *Fusobacterium* is a common taxon found in the gut of canids (Chen et al. 2022; Bragg et al. 2020; Alessandri et al. 2019) and has been linked to protein-rich diets in domestic dogs (Pilla and Suchodolski 2020). Barraza-Guerrero et al. (2021) reported higher abundances of *Fusobacterium* in Mexican wolves (*Canis lupus baileyi*) fed a fresh, raw meat-based diet compared to wolves fed a commercial dry dog kibble-based diet. It is possible that this genus has a similar role in the gut of maned wolves and led to the negative relationship reported between fiber GE and *Fusobacterium* in the present study. Interestingly, a decrease in this genus has also been associated with GI disease in domestic dogs and humans (Scarsella et al. 2023; Félix et al. 2022; Pilla and Suchodolski 2020). *Fusobacterium* spp. are able to metabolize amino acids and can produce the SCFA butyrate, which in general supports GI health, underscoring the role that this genus could play in maintaining GI health (Shah et al. 2024; Félix et al. 2022). This highlights the variable role of *Fusobacterium* in the gut of canids and the need for additional research to assess the relationship between this genus, diet and GI health in the maned wolf specifically.

Zoo-managed maned wolves living alone had higher FA1PI concentrations compared to wolves housed as a breeding pair or same sex sibling group. In general, the maned wolf is monogamous with breeding pairs sharing resources and territory during breeding season and if juveniles are involved (Emmons 2012; Silveira et al. 2009). Stress, including both environmental and psychological, can have a negative influence on intestinal mucosa inflammation via glucocorticoid and/or oxidative stress production (Fan et al. 2023). When an individual is exposed to a stressor, the body responds by activating the hypothalamic-pituitary-adrenal (HPA) axis and releasing glucocorticoids which can go on to alter gut microbiome composition, intestinal barrier function and/or inflammatory responses (Yang et al. 2022). It is possible that an unaccounted-for stressor, factors like temperature, humidity, exposure to novel items, inability to perform natural behaviors, inadequate space allowance and/or restraint (Fan et al. 2023; Yang et al. 2022), could induce an inflammatory response in the intestinal mucosa. This could lead to increased FA1PI concentrations seen in wolves living alone compared to wolves living in a breeding pair or same sex sibling group. Also, it is important to highlight the uneven distribution between social groups (alone n = 17, breeding pair n = 5, same sex sibling group n = 9) which could have influenced our results as well.

We found no relationship between SCFA peak heights or genetic relatedness and gut microbiome composition. Previous literature has established that gut microbiota ferment dietary carbohydrates and create SCFAs as by-products, but bacteria can also use protein and additional dietary components to produce SCFAs (Pilla and Suchodolski 2020; Ríos-Covián et al. 2016; Vital et al. 2015). We did report a negative relationship between fiber GE and abundance of *Fusobacterium* sp., a known butyrate producer (Shah et al. 2024; Félix et al. 2022). However, this genus only represented approximately 0.9% of overall average bacterial abundance. As such, it is not likely that the reported changes in *Fusobacterium* abundance impacted the overall SCFA composition, and thus, leading to the lack of relationship between gut microbiome composition and SCFA peak heights in the present study. Likewise, previous studies have provided evidence that genetically related individuals have more similar gut microbiome composition compared to individuals not genetically related (Cheng et al. 2024; DeCandia et al. 2021). Results from the current study do not support a link between genetic relatedness and gut microbiome composition; it could be due to the stronger influence of the environment on microbiome composition (Bensch et al. 2023; Rothschild et al. 2018), the small sample size for this analysis (n = 22) or both.

While no relationship was documented between gut bacteria and overall SCFA profiles, we identified links between one ASV and propanoic acid as well as multiple ASVs and butanoic and pentanoic acid in zoo managed maned wolves. A negative relationship was identified between *Blautia* sp. (ASV11) and propanoic acid. Among the 14 ASV linked to butanoic acid, we highlight a positive relationship between butanoic acid and *Blautia* sp. (ASV22). Similarly, among the 28 ASVs linked to pentanoic acid, we highlighted another *Blautia* sp. (ASV15) that had a negative relationship while *Megamonas* sp. (ASV15) had a positive relationship with pentantoic acid. The genus *Blautia* has been associated with GI health and inflammation in dogs, with dogs suffering from inflammatory bowel disease and chronic diarrhea having decreased abundance of these bacteria (AlShawaqfeh et al. 2017). Further, zoo-managed red wolves (*Canis rufus*) with loose stool consistency, a proxy for GI health, had an increased abundance of *Blautia* sp. compared to wolves with normal stool consistency (Bragg et al. 2020). Previous literature highlighted a positive relationship between *Blautia hansenii*, *Blautia coccoides* and pentanoic acid in dog DSS-induced colitis systems (Zhang et al. 2025) and mice fed a low-fiber diet supplemented with *B. coccoides* (Holmberg et al. 2024). While the patterns between *Blautia* sp. and pentanoic acid observed in this study are opposite to results from previous literature, our current findings have further linked *Blautia* sp. to gut health modulation via SCFA production (Abdugheni et al. 2022). We also observed that pentanoic acid had a positive relationship with *Megamonas* sp. Previously, *Megamonas* sp. has been associated with acetic and propanoic acid production (Sakon et al. 2008); however, there was a lack of information linking *Megamonas* sp. and pentanoic acid. *Clostridium sensu stricto* sp. (ASV75) but a positive relationship with *Faecalimonas* sp. (ASV47). The genus *Clostridium sensu stricto* can produce acetate, propionate, and butyrate via carbohydrate fermentation that supports GI health (Li et al. 2023). However, this genus has also been identified as an opportunistic pathogen and associated with GI inflammation and enteropathy in domestic dogs (Doulidis et al. 2023; Honneffer et al. 2014). Further research is required to fully elucidate these relationships in this species, but these results have provided potential microbial targets for future research focused on managing GI inflammation in zoo-managed maned wolves.

Utilizing structural equation modeling, we determined that fiber GE had separate direct, negative effects of fiber GE on *Fusobacterium* sp. abundance and on FA1PI concentrations but there were no indirect effects of *Fusobacterium* sp. on FA1PI concentrations. This means that dietary fiber GE directly impacted *Fusobacterium* sp. abundance and FA1PI concentrations, but the direct effects of fiber GE are separate and there is no indirect modulation of FA1PI concentrations by changes in *Fusobacterium* sp. abundance caused by fiber GE. While previous literature does not directly link dietary fiber and the genus *Fusobacterium*, there has been a relationship established between dietary protein and increased *Fusobacterium* abundance and dietary fiber and decreased *Fusobacterium* abundance in canids (Bermingham et al. 2017; Butowski et al. 2022). Therefore, result from the current study indicate that wolves provided with a diet with high fiber GE had a decrease in the abundance of *Fusobacterium* sp. This pattern has been previously documented in canids (Panasevich et al. 2015; Kim et al. 2017; Sandri et al. 2017; Schmidt et al. 2018). The negative, direct effect of fiber GE on FA1PI concentrations could be a result of the general benefit to GI health by including fiber in the diet (Montserrat-Malagarriga et al. 2024; Cave et al. 2023), due to fiber increasing gut transit time, having a laxative effect and shortening the time for FA1PI concentrations to accumulate in the feces (Müller et al. 2018), or both.

## 5. Conclusion

This study was the first to characterize the gut microbiome of zoo-managed maned wolves and examine the relationship among gut bacteria absolute abundance and environmental factors, genetic relatedness and SCFA composition in this species. Our results underscore how management choices in zoological facilities can influence the gut bacterial community and possibly GI health. Specifically, we encourage future research on fiber intake, *Blatuia* sp. abundance, SCFA concentrations, and GI health to fully elucidate how diet management choices and/or mediation with specific bacteria linked to SCFAs could be used as a tool to support GI health in zoo-managed maned wolves. The taxonomic uniqueness and lack of knowledge about this species creates a challenge for zoological facilities. Wolves that have poor GI health can also suffer from reduced reproductive fitness (McKenzie et al. 2017), a main priority for a vulnerable, zoo-managed species. Maintaining a healthy zoo-managed population of maned wolves is vital to ensure the continued existence of this unique and vulnerable canid.

## CRediT authorship contribution statement

**Morgan Bragg**: Conceptualization, Methodology, Formal analysis, Investigation, Data curation, Writing – Original draft, Visualization. **Carly R. Muletz-Wolz**: Conceptualization, Methodology, Formal analysis, Resources, Writing – Review & Editing, Supervision, Funding acquisition. **Elizabeth W. Freeman**: Resources, Writing – Review & Editing, Supervision. **Nucharin Songsasen**: Conceptualization, Methodology, Resources, Writing – Review & Editing, Supervision, Funding acquisition.

## Data Accessibility

The datasets presented in this study can be found on the National Center for Biotechnology Institute – Sequence Read Archive. The repository/repositories and accession number(s) can be found below: https://www.ncbi.nlm.nih.gov/PRJNA1450688. Code and metadata files can be found at: https://github.com/morganbragg/ManedWolfGutMicrobiome

## Ethics statement

The animal study was reviewed and approved by the IACUC—George Mason University and IACUC—Smithsonian Institution.

## Acknowledgements

We would like to thank all the facilities that participated in this study; Fossil Rim Wildlife Center, Smithsonian Conservation Biological Institute, Wildlife Safari, Sedgwick County Zoo, White Oak Conservation, and Little Rock Zoo. This work was funded by the Morris Animal Foundation [grant ID D19ZO-062].

